# Automated collection of pathogen-specific diagnostic data for real-time syndromic epidemiological studies

**DOI:** 10.1101/157156

**Authors:** Lindsay Meyers, Christine C. Ginocchio, Aimie N. Faucett, Frederick S. Nolte, Per H. Gesteland, Amy Leber, Diane Janowiak, Virginia Donovan, Jennifer Dien Bard, Silvia Spitzer, Kathleen A. Stellrecht, Hossein Salimnia, Rangaraj Selvarangan, Stefan Juretschko, Judy A. Daly, Jeremy C. Wallentine, Kristy Lindsey, Franklin Moore, Sharon L. Reed, Maria Aguero-Rosenfeld, Paul D. Fey, Gregory A. Storch, Steve J. Melnick, Christine C. Robinson, Jennifer F. Meredith, Camille V. Cook, Robert K. Nelson, Jay D. Jones, Samuel V. Scarpino, Benjamin M. Althouse, Kirk M. Ririe, Bradley A. Malin, Mark A. Poritz

## Abstract

Health-care and public health professionals rely on accurate, real-time monitoring of infectious diseases for outbreak preparedness and response. Early detection of outbreaks is improved by systems that are pathogen-specific. We describe a system, FilmArray^®^ Trend, for rapid disease reporting that is syndrome-based but pathogen-specific. Results from a multiplex molecular diagnostic test are sent directly to a cloud database. www.syndromictrends.com presents these data in near real-time. Trend preserves patient privacy by removing or obfuscating patient identifiers. We summarize the respiratory pathogen results, for 20 organisms from 344,000 patient samples acquired as standard of care testing over the last four years from 20 clinical laboratories in the United States. The majority of pathogens show influenza-like seasonality, rhinovirus has fall and spring peaks and adenovirus and bacterial pathogens show constant detection over the year. Interestingly, the rate of pathogen co-detections, on average 7.7%, matches predictions based on the relative abundance of organisms present.

## Introduction

The availability of real-time surveillance data that can monitor the spread of infectious diseases benefits public health [1-3]. At present, tracking of respiratory or foodborne outbreaks relies on a variety of methods ranging from automated real-time electronic reporting to manual web entry of test results. Systems such as the Centers for Disease Control and Prevention’s (CDC) FluView [4, 5], National Respiratory and Enteric Virus Surveillance Systems (NREVSS, [6]), National Electronic Disease Surveillance System (NEDSS [7, 8]), Global Emerging Infections Surveillance (GEIS) [9], and others, although web-based still require manual entry of data from laboratories, resulting in data that are often incomplete or not current.

Newer efforts at syndrome-based surveillance [10-12] include BioSense (extraction of symptomatic data from electronic health records [13]), Google Flu (tracking of Internet search queries [14] but recently discontinued [15]), and Flu Near You (voluntary reporting [16]). Additionally, numerous next generation, syndromic surveillance systems; e.g., pharmacy sales records [17, 18], Twitter conversations [19, 20], and Wikipedia hits [21, 22], have come online in the past five years. However, these systems cannot report the specific pathogen causing an increase in a particular set of symptoms. Finally, there are more localized efforts such as GermWatch in Utah [23, 24] and the Electronic Clinical Laboratory Reporting System (ECLRS) in New York [25, 26] that draw from hospital (HIS) and laboratory (LIS) information systems. This disparity in technologies and data collection methods results in incomplete surveillance.

Comprehensive and uniform diagnostic test data will aid in the identification of potential outbreaks. A combination of broad respiratory pathogen testing and an internal electronic surveillance system enabled the rapid dissemination of data across the largest health care system in New York, the North Shore-LIJ Health System (now Northwell Health) during the influenza A H1N1-2009 pandemic in the New York City area. Pathogen-specific molecular testing permitted rapid a) notification to state epidemiologists; b) tracking of the virus so that health care resources could be managed effectively; and c) evaluation of influenza diagnostics [27, 28]. Today, with the threat of emerging pathogens such as Middle East Respiratory Syndrome coronavirus (MERS CoV), avian influenza, enterovirus D68, and Ebola virus, real-time surveillance programs are critical [1, 29, 30].

It is not always possible to accurately diagnose the causative agents of most infectious diseases from symptoms alone due to overlapping clinical presentation. Thus, to achieve maximal utility, infectious disease surveillance systems should move beyond syndrome-based reporting and be pathogen-specific and comprehensive, reporting on as many of the common pathogens for a particular syndrome as possible. Sensitive and specific automated molecular diagnostic systems that detect up to four different pathogens in a single sample have been available from in vitro diagnostic (IVD) manufacturers for some time [31-33]. However adoption of IVD platforms with broad multiplexing capability has become widespread only in the last few years. Commercially available systems that can detect most of the known etiological agents for respiratory, gastrointestinal and other multi-pathogen syndromes [34-36] include the BioFire (Salt Lake City, UT) FilmArray^®^ System [37]; the GenMark (Carlsbad, CA) eSensor XT-8^®^ [38] and ePlex^®^ [39]; and the Luminex (Austin, TX) xTAG^®^ [40], nxTag^®^ [41] and Verigene^®^ systems [42].

Multi-analyte diagnostic tests provide the raw data needed for real-time pathogen-specific surveillance but there remain a number of obstacles to sharing these results (reviewed in [43]). The obstacles largely center on information privacy and network security. A real-time surveillance system using diagnostic test results requires safeguards for protected health information (PHI). Medical records and devices have become attractive targets for cyber attackers in recent years [44], which has made hospitals and clinics reluctant to connect their Local Area Networks (LANs) to the Internet. Releasing patient test results requires the removal of PHI or authorization from the patient. Studies have shown that de-identification of patient data is not as simple as removing all specific identifiers because, in the age of big data, under the right circumstances it is possible to re-associate patients and their data using publicly available information [45-48].

We describe here the implementation of a real-time pathogen-specific surveillance system that overcomes the PHI concerns noted above. FilmArray Trend de-identifies, aggregates and exports test results from FilmArray Instruments in use in US clinical laboratories. Although data from all commercially available FilmArray Panels (Methods and [49]) is exported to the Trend database, we focus here on the Respiratory Panel (RP), which can detect 17 viral (adenovirus [ADV], coronaviruses [CoV OC43, 229E, NL63, HKU-1], human metapneumovirus [hMPV], human rhinovirus/enterovirus [HRV/EV]; influenza A [Flu A, subtyping H1N1, 2009 H1N1, H3N2], influenza B [Flu B]; parainfluenza viruses [PIV1-4], and respiratory syncytial virus [RSV]), and three bacterial (*Bordetella pertussis, Chlamydia pneumoniae, and Mycoplasma pneumoniae*) pathogens [37, 50-56]

With more than 344,000 patient results for the FilmArray RP test alone, the Trend database has many of the properties associated with “big data” as it applies to infectious disease [57]. After describing how the dataset can be cleaned of non-patient tests, we make some observations on the seasonality of the different respiratory pathogens and we apply the ecological concept of “species diversity” [58] to observe a correlation between the abundance of each pathogen and the rate at which co-detections (more than one positive result per test) occur.

## Results

### Sending FilmArray data directly to the cloud

The FilmArray Trend public website is an outgrowth of the Trend clinical website, which was developed to provide clinical laboratories using the FilmArray System with up-to-date information on the respiratory, gastrointestinal and meningitis/encephalitis pathogens circulating at their institutions. The most general and efficient way to export both the clinical and public Trend information is to follow a “Bottom-Out” approach to data export (Figure 1). In this scheme, the FilmArray Instrument sends data via the Internet directly to a single cloud database where it can be viewed by health care providers at the originating institution. This data export pathway contrasts with a “Top-Out” approach (Figure 1) in which diagnostic test results are pushed from the instrument up to the LIS, then to the HIS and, finally, a subset of this information is forwarded to cloud-based databases.

**Figure 1.**
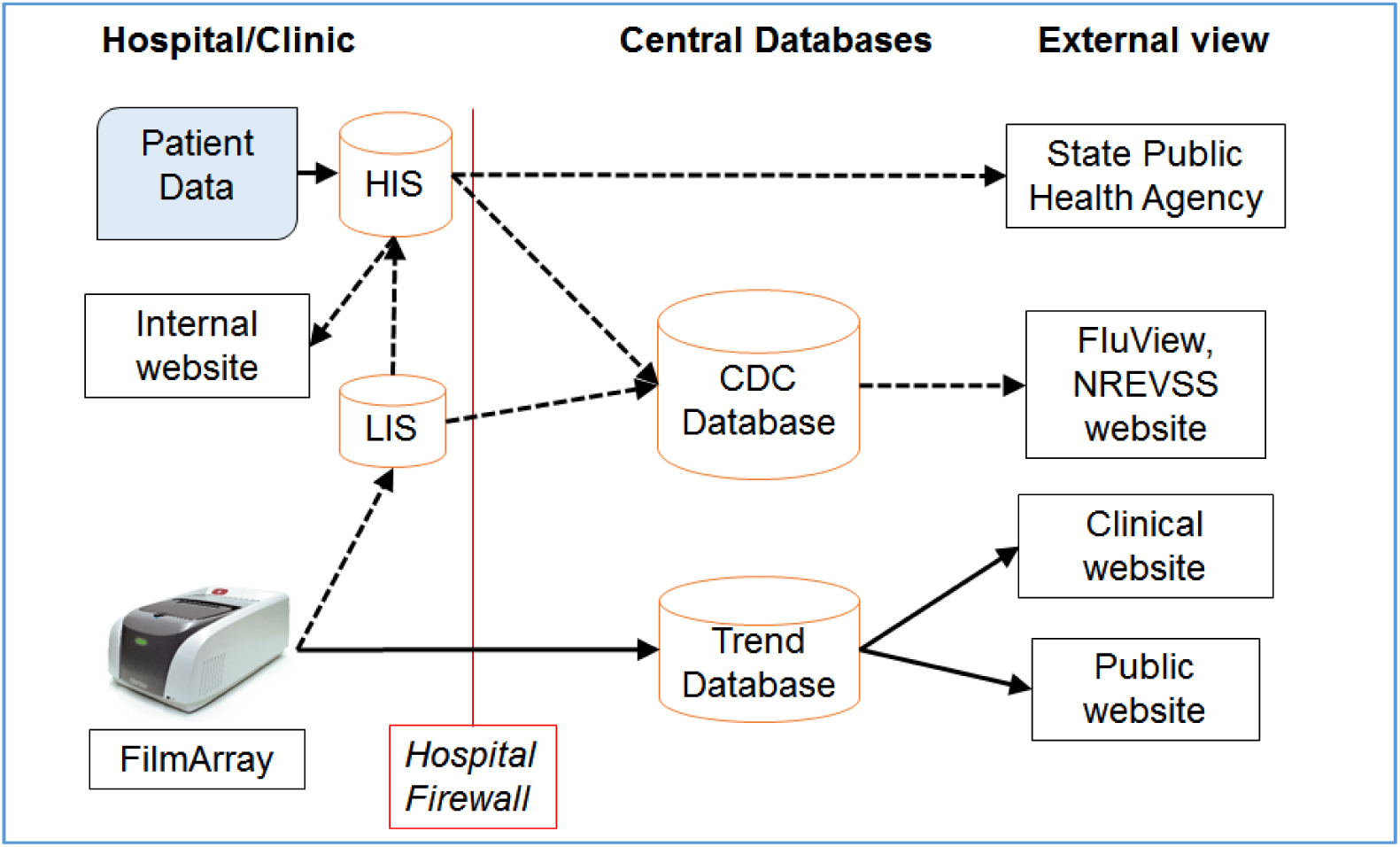
Schema for export of IVD test results to an external database. Bottom-Out and Top-Out approaches for data export are indicated by solid and dashed lines respectively. Some institutions have developed their own systems for aggregating and displaying infectious disease data (indicated by “Internal website”). Other abbreviations are defined in the text.

Initial testing of the Trend export mechanism was performed in collaboration with the Clinical Laboratories of the Medical University of South Carolina. This trial allowed us to develop and test auto-export functions and de-identification protocols for the Trend software. The de-identification requirement of the Health Insurance Portability and Accountability Act (HIPAA) of 1996 [59], specifically the Safe Harbor provision, requires the removal of 18 enumerated variables that could directly or indirectly identify an individual. In accord with this requirement, the first stage study did not export test identifiers, or free form text fields and only returned the year of the test. The initial dataset provided low-resolution information but was a useful platform to evaluate the proposed system. Further development to enable export of higher resolution data required the design of routines that would adhere to an alternative HIPAA de-identification strategy, namely, the Expert Determination approach, which requires a risk assessment demonstrating that the chance of re-identifying an individual is sufficiently small [60, 61]. The Expert Determination process identified and made recommendations for fields that could facilitate disclosure of PHI. A summary of the Expert Determination results detailing the risk of Trend data in regard to replicability, availability and distinguishability is shown in Methods, Table 2.

**Table 1:** Distribution of number of positive results per test for Trend RP tests.

| # Positives per RP Test | All | Tests after event removal | Final Trend Dataset |
| --- | --- | --- | --- |
| 0 | 48.16 | 48.47 | 48.91 |
| 1 | 42.65 | 43.04 | 43.43 |
| 2 | 6.64 | 6.65 | 6.71 |
| 3 | 0.96 | 0.94 | 0.95 |
| 4 | 0.19 | 0.16 | 0.00 |
| 5 | 0.07 | 0.06 | 0.00 |
| 6 | 0.06 | 0.04 | 0.00 |
| 7 | 0.54 | 0.26 | 0.00 |
| 8 | 0.01 | 0.01 | 0.00 |
| 9 | 0.06 | 0.03 | 0.00 |
| 10 | 0.03 | 0.02 | 0.00 |
| 11 | 0.03 | 0.02 | 0.00 |
| 12 | 0.50 | 0.26 | 0.00 |
| 13 | 0.00 | 0.00 | 0.00 |
| 14 | 0.01 | 0.00 | 0.00 |
| 15 | 0.01 | 0.01 | 0.00 |
| 16 | 0.03 | 0.02 | 0.00 |
| 17 | 0.03 | 0.01 | 0.00 |
| 18 | 0.02 | 0.00 | 0.00 |
| 19 | 0.00 | 0.00 | 0.00 |

Percent of FilmArray RP tests in the Trend dataset that are positive for the indicated number of organisms for all data in the Trend dataset (All), after removing known control tests (Tests after event removal), and what is analyzed here (Final Trend Dataset). Data are scaled to sum to 100%. The panel tests for 20 organisms but a positive for Flu A H1-2009 is reported as negative for Flu A H1 so the maximum possible number of positive results is 19.

**Table 2:** Qualitative analysis for the Expert Determination.

| Field | Replicability | Availability | Action |
| --- | --- | --- | --- |
| Sample | **High** (continually associated with a patient) | **High** (name exists in public records) | Deleted, the Trend database is not configured for the capture of this field. |
| Test Time and Date | **Medium** (only associated with patient once) | **Medium** (may be self-disclosed in public) | Binned into date range when minimum threshold of individuals is met |
| FilmArray Pouch Serial Number | **Medium** (only associated with patient once) | **Low** (only available to institution) | Truncated (last digit removed) |
| Test Location | **High** (likely continually associated with a patient) | **Medium** (may be self-disclosed in public) | Limited to sites drawing from > 20,000 pop. |
| Instrument Name | **Low** (not associated with patient) | **Low** (only available to institution) | No Action |

Because the Expert Determination study established that no PHI will be disclosed by the Trend data export algorithm, Data Use Agreements (DUA), rather than Business Associates Agreements (BAA, see Methods for the difference between a DUA and a BAA) were executed with each of the collaborating institutions (listed in Table 3). The DUAs define for the clinical laboratory how BioFire will manage and make use of the Trend data. The Trend client software residing on the FilmArray computer queries the FilmArray test result local database and exports the results to an Amazon Web Services (AWS) database (see Methods). The Trend client software performs de-identification on the FilmArray computer prior to export as detailed in Table 2. Health care providers are granted access to their institutions Trend data by the Laboratory Director. Since web access to view the data is restricted to the local site, de-identification of geographic indicators is not required. However, in implementation of the public Trend website, which presents the national syndromic test surveillance (Figure 1), we have further aggregated the data with respect to geographic origin and obfuscated the date of the test (see Methods). Since only de-identified data are exported from the clinical institutions, no PHI is sent to or stored on the cloud server.

**Table 3:** Clinical laboratories participating in the initial Trend study.

| HHS region | Name | City, State | Hospital, Facility Type | Bed Count |
| --- | --- | --- | --- | --- |
| 1 | University of Massachusetts Medical School-Baystate | Springfield, MA | University Affiliated | 1,000 |
| 2 | Albany Medical Center | Albany, NY | Medical College Affiliated | 800 |
| 2 | Northwell Health | Lake Success, NY | Network | 6,400 |
| 2 | NYU Langone Health | New York, NY | Network | 1,500 |
| 2 | Stony Brook University Hospital | Stony Brook, NY | University Affiliated | 600 |
| 2 | Winthrop University Hospital | Mineola, NY | University Affiliated | 600 |
| 4 | Greenville Health System | Greenville, SC | Network | 1,400 |
| 4 | Nicklaus Children's Hospital | Miami, FL | Children's | 300 |
| 4 | Medical University of South Carolina | Charleston, SC | University Affiliated | 700 |
| 5 | South Bend Medical Foundation | South Bend, IN | Commercial lab serving regional hospitals | NA |
| 5 | Detroit Medical Center | Detroit, MI | Network | 2,000 |
| 5 | Nationwide Children's Hospital | Columbus, OH | Children's | 600 |
| 7 | Children's Mercy | Kansas City, MO | Children's | 400 |
| 7 | Washington University Medical Center | St. Louis, MO | University Network | 1,700 |
| 7 | Nebraska Medical Center | Omaha, NE | University Network | 600 |
| 8 | Children's Hospital Colorado | Denver, CO | Children's Hospital | 500 |
| 8 | Intermountain Medical Center | Salt Lake City, UT | University Affiliated | 500 |
| 8 | Primary Children's Hospital | Salt Lake City, UT | Children's | 300 |
| 9 | Children's Hospital of Los Angeles | Los Angeles, CA | Children's, University Affiliated | 500 |
| 9 | UC San Diego Health | San Diego, CA | University Affiliated | 700 |

All sites submitted the Trend project for review by their local Institutional Review Board (IRB); all but one of the 20 IRBs deemed the project “exempt” because of the absence of PHI export. Thus the security requirements for the database and the controls necessary for storage and transport of de-identified data are significantly reduced.

Following the protocol established by Expert Determination review, the Trend software delays the export of data until the number of tests queued for export exceeds a minimum threshold for each type of FilmArray panel. In practice, this results in an average time to export of less than two hours from each site that has multiple instruments. 98.5% of the tests are exported within 24 hours.

### Characteristics of the FilmArray sites used in the Trend pilot study

The 20 sites contributing to the Trend pilot project (Methods, Table 3) have the same average number of instruments, six (range one to 22) as for all US FilmArray customers. The Trend pilot sites have been using the FilmArray RP test for an average of 3.8 years (range one to six) prior to July 2017. The size of the institutions participating ranges from 300 to 6400 beds with the majority being large hospitals and health care networks with an average of 1,100 beds. Six (30%) sites are pediatric hospitals and one is a reference laboratory. Fifteen (75%) of the sites have uploaded archived FilmArray RP test results to the Trend database, with eight (40%) reporting results dating back to 2012. Unless stated otherwise, the data presented here covers the period from July, 2013 to July, 2017.

The algorithm used to diagnose the cause of respiratory disease varies by site. More than half of the Trend sites do not enforce an institutional respiratory testing protocol and, even within sites that have a required protocol, some discretionary use of FilmArray RP is allowed. Without detailed records from each institution’s HIS, it is not possible to determine whether the FilmArray RP was used as a front line test or as a reflex test (typically following a negative result for influenza and/or RSV).

### Cleaning non patient test results from the FilmArray Trend database

To determine the prevalence of respiratory pathogens, we needed to expunge the Trend database of test results that are not derived from clinical patient samples. Non-patient results come from a variety of sources including verification testing, routine Quality Control (QC) and proficiency testing (PT) (see Methods, Cleaning the Trend data). Despite this complexity, the majority of non-patient test results can be identified and distinguished from the patient-derived data because of the high number of positive organism calls in a single test and because of the temporal aspects of verification and control testing (see Methods). QC tests are estimated to account for half of all FilmArray RP results in which more than 3 organisms are detected. A combination of filters (Methods) removes 3.52% of the total tests from the Trend data set. We estimate that 3% of the remaining dual and 6% of the remaining triple detection tests respectively may come from QC testing. In addition to the exclusion of tests temporally associated with validation events, all results with four or more positives were removed from further analysis (0.9% of the filtered total total). This includes the 0.16% of test results with exactly four organisms (Table 1, “Tests after event removal” column) because the minority are derived from patient testing.

### Detection of respiratory pathogens in Trend samples from 2013 to 2017

The detection counts and pathogen detection rates derived from the Trend data set for each organism in the FilmArray RP are shown in Figure 2. Other views of these data, including percent detection of individual organisms or combinations of organisms, are available on the FilmArray Trend website: www.syndromictrends.com. The FilmArray RP Test Utilization Rate (RP TUR, see Methods) and the individual organism detection counts increased over this period because the Trend clinical sites increased their utilization of the FilmArray RP tests (Figure 2A). Seasonal fluctuations can also be seen within this growth pattern, with use increasing up to fourfold each winter when compared to the previous summer. HRV/EV, the most common pathogen detected group, is identified in approximately 30% of all samples tested each year (Figure 2-figure supplement 1). Other pathogens detected in five to 10% of the samples include: RSV, the PIVs, ADV, Flu A and hMPV. *M. pneumoniae, C. pneumoniae*, *B. pertussis* and Flu B are detected in less than 2% of all samples. The average percentage of each organism is relatively constant over the four years of data in the Trend database (Figure 2-figure supplement 2).

**Figure 2.**
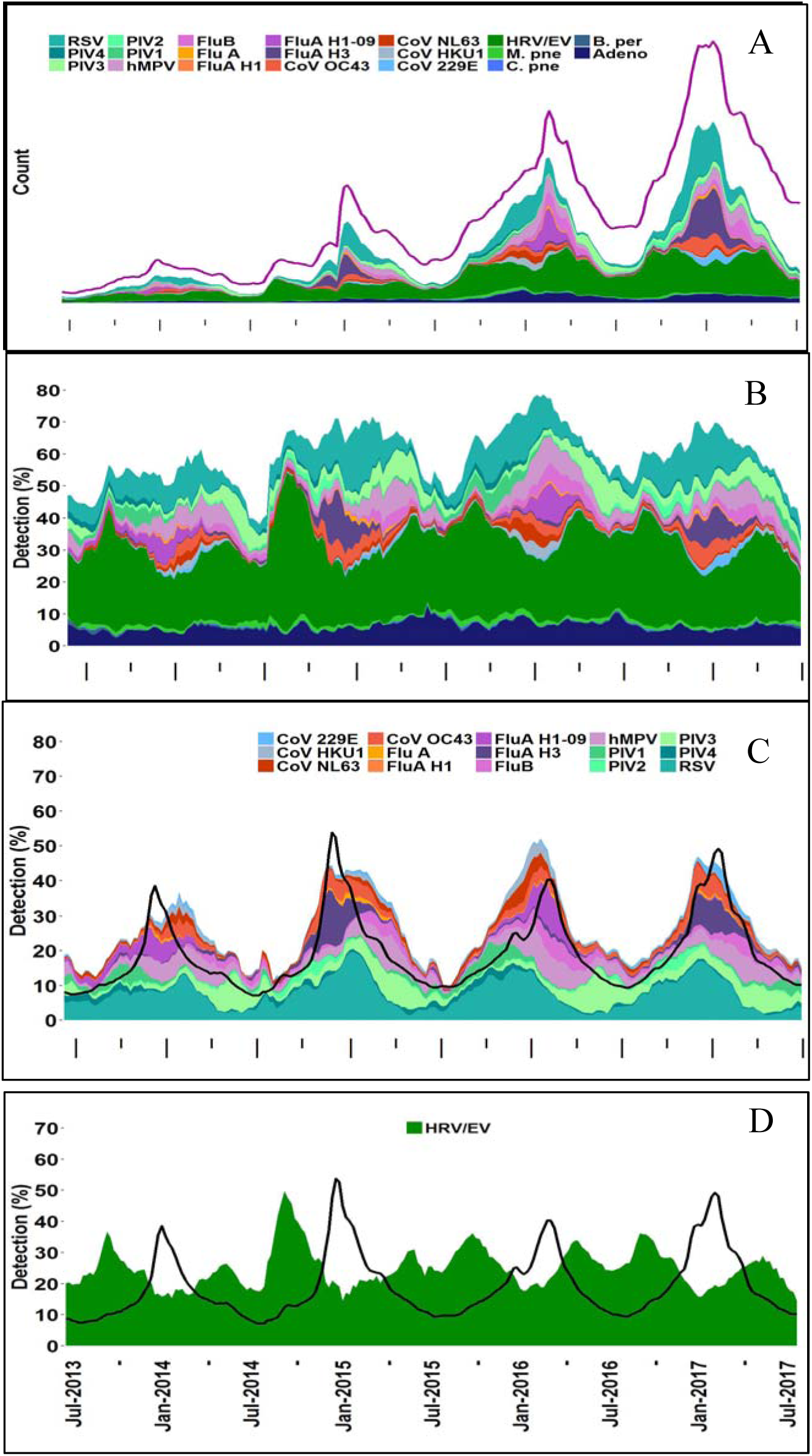
Detection of RP organisms over time across all sites. **Figure 2 legend:** Detection of FilmArray RP pathogens in the Trend dataset displayed as a stacked area graphs. All panels have the same time period (July 2013 to June 2017). A) Count of each organism. The TUR metric (red line, units are FilmArray RP tests performed), and count of FilmArray RP tests that are negative (white area between pathogen count and TUR) are indicated. The y-axis values are not indicated as this is considered proprietary information. B) Pathogen detection rates for all organisms. C) Pathogen detection rates for the subset of organisms that show seasonality (see Results and the legend for the list of organisms). D) HRV/EV detection rates. CDC measured ILI is indicated (black line) in panels C and D. Organisms follow the same color scheme in all panels; the order of organisms in the legend (down then across) matches the stacked area graph top to bottom. **Figure 2-Figure Supplement 1:**Detection of adenovirus and three bacteria **Figure 2-Figure Supplement 2**:Detection of FilmArray RP organisms by type **Figure 2-Figure Supplement 3**:Organism detection by type and year

The pathogens’ seasonal variability measured by percent detection can be classified into at least three groups. **Group 1**: The majority of organisms follow the classical “respiratory” season (October through March) and increase by more than 10-fold above their baseline detection rate (Figure 2C). These include the CoVs, Flu A, Flu B, hMPV, the PIVs, and RSV (PIV3 is a slight exception to this rule in that it peaks in the summer months, and has a winter peak that is only detected regionally (data not shown)). Within this group, all but five viruses demonstrate significant biennial fluctuations; Flu B, hMPV, OC43, and PIV3 and RSV experience relatively consistent annual peaks. **Group 2**: HRV/EV is in a class by itself in that it is detected in a high percentage of tests over time, (minimum of 10% in winter) and experiences moderate peaks of two to three fold outside the respiratory season baseline, in the early fall and spring (Figure 2D). **Group 3**: The bacteria and adenovirus are present at a relatively constant rate (Figure 2-Figure supplement 1). The CDC FluView reported rate of Influenza-Like-Illness (ILI) tracks moderately well with the Group 1 organisms (cross correlation of 0.85) and not with HRV/EV or with ADV and the bacteria.

### Comparison of Trend to CDC measures of influenza

The CDC FluView network [4, 5] gathers information about influenza prevalence from a large number of public health and clinical laboratories in the US. FluView is considered the gold standard for these measures. We compared the Trend detection rates for Flu A (all subtypes) plus Flu B to the FluView Influenza (A and B) from September 2015 to July 2017 (Figure 3). The analysis was restricted to this time period because of a change in the CDC’s reporting of Flu prevalence in the fall of 2015. A cross-correlation of 0.974 was observed between the Trend Flu A/B percent detection and FluView reported influenza prevalence. Notably, the onset, peak and duration of the influenza season coincide between the two measures.

**Figure 3:**
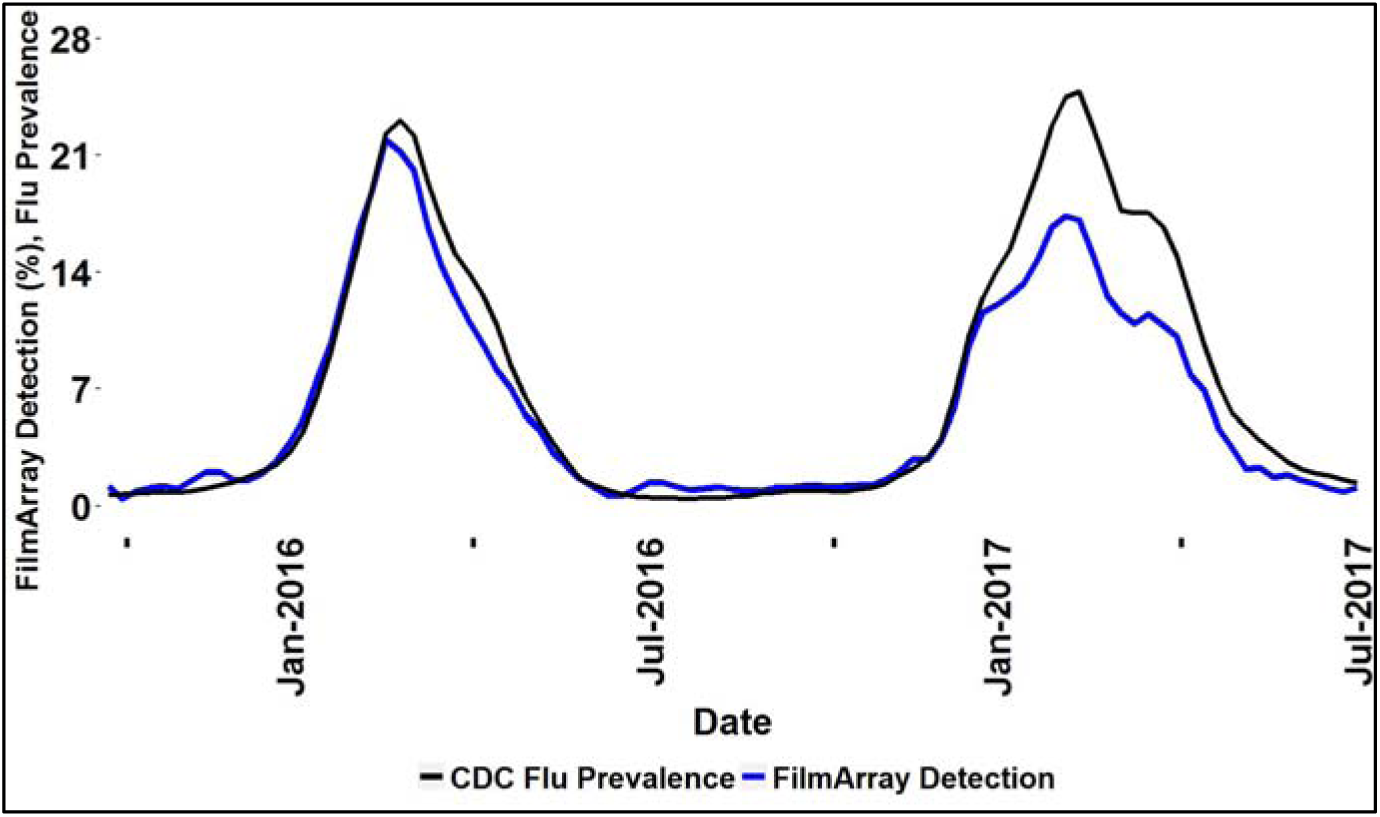
Trend Influenza Detection Rate compared to CDC Influenza activity. Percent of combined FilmArray Flu A (all subtypes) and Flu B detections (blue line) and CDC reported Influenza prevalence (black lines). CDC data are aggregated only from regions with participating Trend sites.

### Respiratory Panel Co Detections

We found that 26,347 FilmArray RP tests in the Trend dataset had 2 or 3 co-detections (7.7% of total FilmArray RP tests) and that the co-detection rate of each organism varies widely (10-55%). Although an additional pathogen was detected in half of the ADV and CoV positive samples, co-detections were observed in only 10-15% of the samples positive for either Flu A or Flu B (Figure 4).

**Figure 4:**
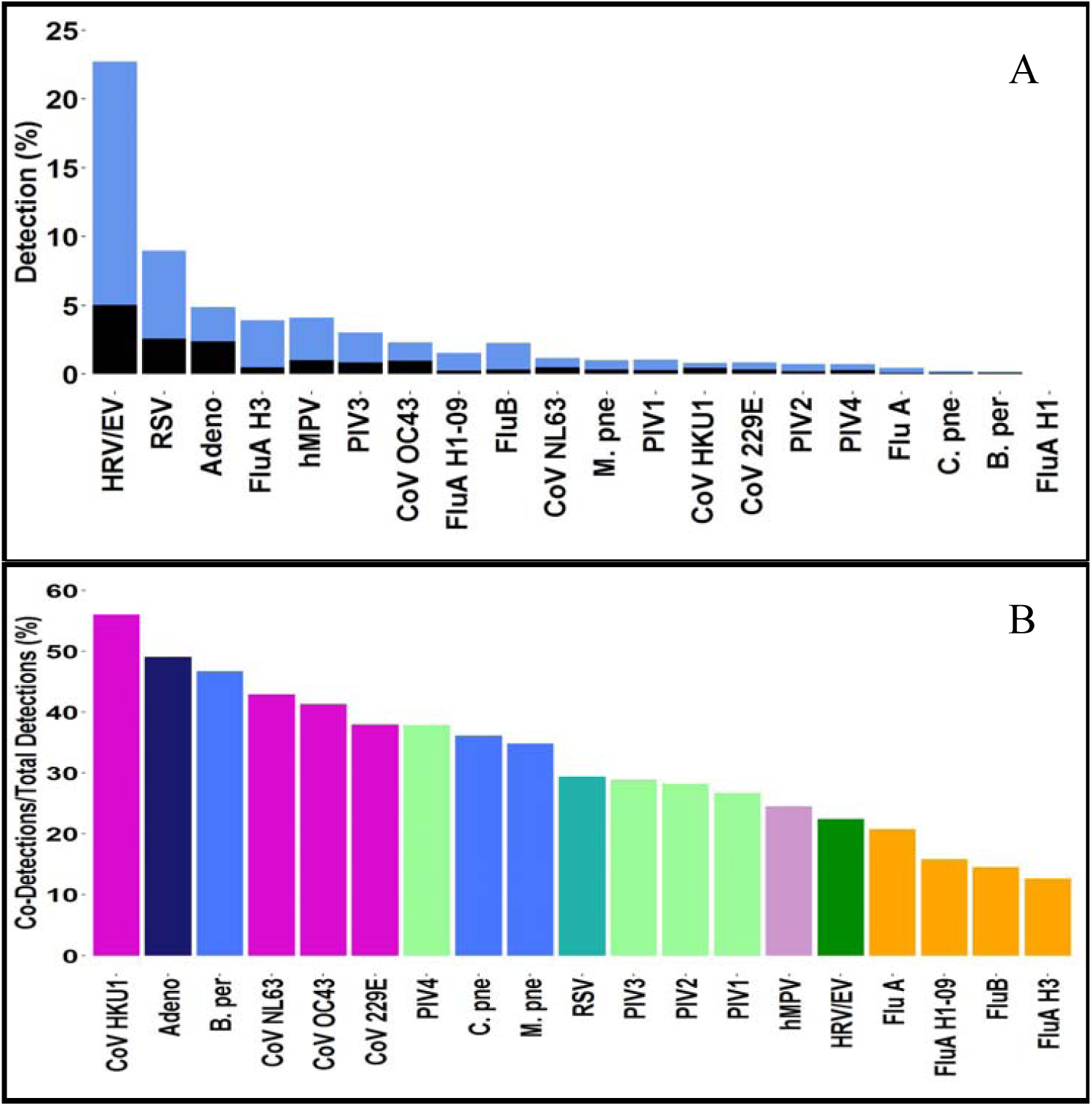
Detection rates for all organisms compared to co-detections. **(A)** Percent total positive detections for each organism in the RP Trend data set is presented in stacked bars, showing the rate of detection of a single organism (A, blue) and those involved in a co-detection (A, black). Data are calculated for each site during the period of July 2013 to March 2017, when available, and then aggregated. **(B)** Percentage of each organism involved in a co-detection is shown. Bars are colored by pathogen family (CoV, purple; Bacteria, blue; PIV green; Flu A yellow).

Trend data have high temporal, spatial, and organism-specific resolution. These three properties allow for a novel evaluation of co-detections. The observed rates of co-detections should be influenced by the number of circulating pathogens detected by the FilmArray RP test at a particular site. Figure 5A shows the average number of unique organisms detected at each site in a given week (see Methods: Calculation of co-detection rates). This number fluctuates from a summer low of four to a winter high of 11 pathogens. Figure 5B (grey bars) shows that the total rate of organism co-detections in the Trend dataset fluctuates annually with peak rates occurring in the winter months. The average rates have been as high as 12% in the winter of 2016 and as low as 2% in the summer of 2014.

**Figure 5:**
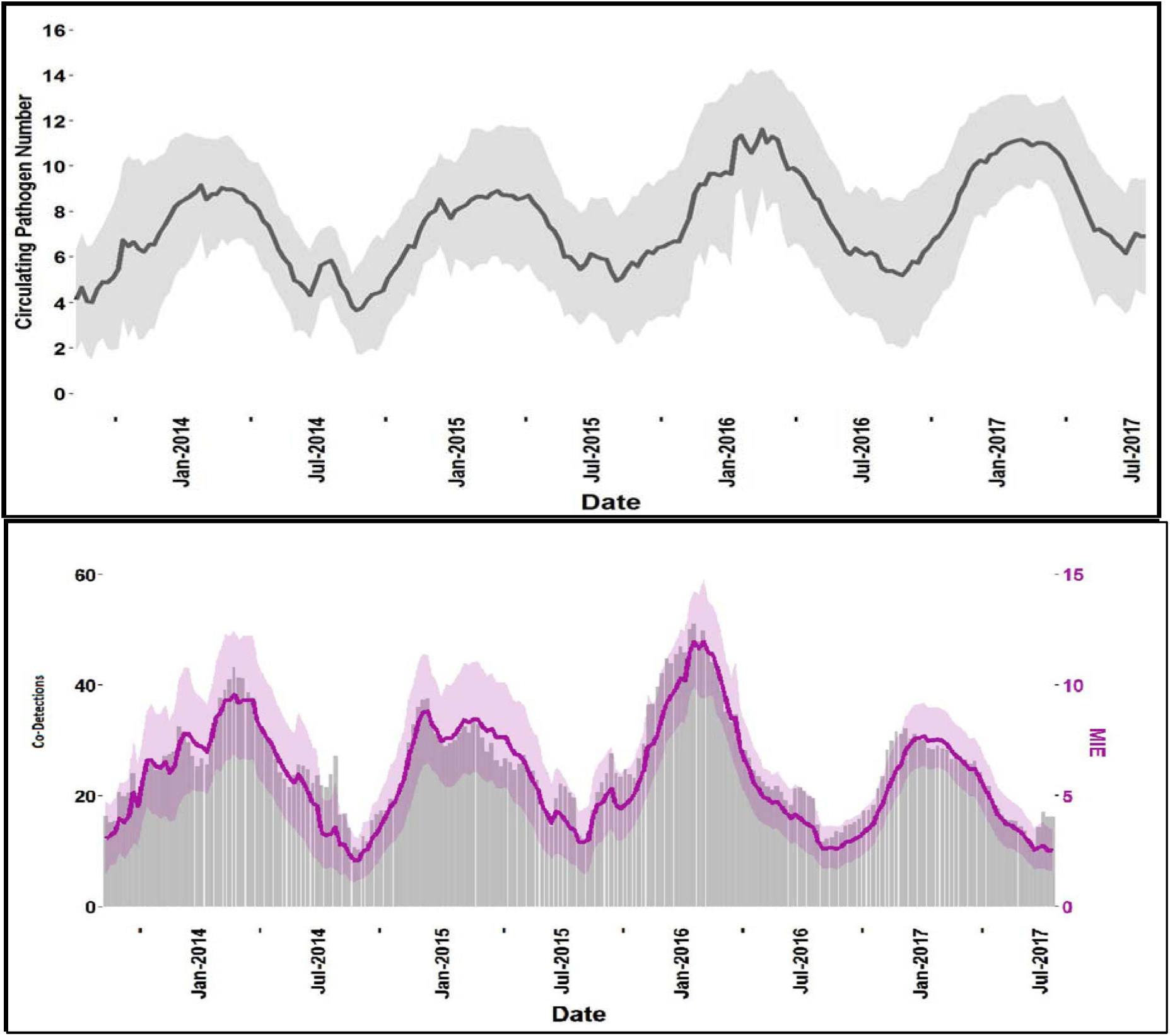
Seasonal variation in pathogen diversity and co-detections. **A)** Average circulating pathogen number (black line) and one standard deviation computed across all Trend sites (grey area). **B)** Rate of co-detections in the RP Trend dataset (grey bars, left axis) the MIE index (purple line, right axis) and MIE confidence intervals (shaded purple area). **Figure 5-Figure supplement 1**: Linear regression of MIE and observed co-detections

From the Trend data, a Measure of Interspecific Encounter (MIE) can be calculated as the probability of a co-detection, weighted by the prevalence of each circulating pathogen at a site. Although the value of the MIE metric is higher than the actual co-detection rate it correlates well (Figure 5B, purple line compared to the grey bars has a cross correlation of 0.9488 at a lag of 0). The magnitude adjustment between MIE and the observed co-detections is calculated by the slope of the linear regression of the two metrics (Figure 5 n figure Supplement 1) and has a value of 4.05 (R^2^ = 0.9003).

## Discussion

This article describes “FilmArray Trend”, a new system for real-time reporting of widespread pathogen-specific syndromic data. This system already has many of the important properties of big data. We consider Trend in terms of the “V”s that are often used to describe big data: volume (amount), velocity (speed of acquisition), veracity (accuracy), variety (diversity of information) and value (utility) [57, 62].

### Trend Volume

The Trend RP dataset is growing at an average rate of >200,000 pathogen test results per month. Connecting the first 20 clinical sites has provided insight into the principal concerns that will be raised by the legal, IT and administrative departments of the healthcare providers that house FilmArray instruments. It should be possible, therefore, to expand the Trend installed base by 10 to 20-fold over the next few years. Similarly, the existence of Trend should enable other IVD manufacturers to build their own Trend-like systems with greater acceptance on the part of their customers, thereby allowing a more global and comprehensive surveillance perspective.

### Trend Velocity

The data in Figure 2 are similar to previous demonstrations of the seasonality associated with different respiratory viruses [63-66]. What is novel is that this data is generated automatically, on site, and in close to real-time compared to other surveillance systems. Greater than 98% of the test results are exported to the Trend database within 24 hours of being generated. As part of the de-identification protocol, sequential FilmArray RP tests of the same type are put into the same time bin. This has the effect that test results are exported faster during periods of peak use such as during the peak of the respiratory season or during an outbreak. Trend should be instrumental at a local level to determine the start of respiratory season; many hospitals make significant changes to their operations based on this event however, at present, data collection to track the respiratory season is often slow and manual, or semi-automated at best.

The key to implementing Trend clinical sites was to demonstrate that FilmArray test results can be exported without the risk of breaching PHI confidentiality either directly or through some combination of the data that was exported. Trend successfully used the Expert Determination process as prescribed by the HIPAA guidelines (see Methods), which greatly simplified the data sharing agreement between BioFire Diagnostics and the clinical site and allowed health care providers to use Trend without risk of inadvertently disclosing PHI.

The software architecture underlying the Trend system is both simple and secure: 1) no changes to the institutional firewall or LAN are needed; 2) the Trend database cannot reach back and query the FilmArray computer due to the institutional firewall, which is set to outbound data only; 3) Trend software can only submit data to the cloud database and cannot query the database. Yet, despite this security, authorized users of the Trend database can mine the deidentified data to look for novel patterns in respiratory pathogen epidemiology.

### Trend Veracity

The goal of an epidemiological surveillance network is to infer which infectious diseases are circulating in the general population based on testing a sample of patients [67]. Different surveillance systems have different biases in their data, biases that perturb the ability to predict true population prevalence.

While the removal of all PHI has great benefits in terms of implementation, it also has several shortcomings that complicate interpretation of the data. First, Trend cannot account for the variability in the diagnostic testing algorithms applied to the selection of samples to be tested by the FilmArray Instruments. During respiratory season, health care providers may prescreen patients with other diagnostic tests including rapid antigen or molecular assays for influenza and RSV, and/or commercial and laboratory developed molecular tests for a mix of other respiratory pathogens. Depending upon the sensitivity of these upstream tests, 50-90% of influenza and RSV for the subset of the patients screened would be excluded from the Trend dataset if the front line test is positive. This testing protocol may skew the actual prevalence of not only influenza and RSV but all other individual respiratory pathogens and co-infections detected by the FilmArray. In some institutions, testing is reserved for hospitalized patients and others at risk for developing complications of respiratory tract infections including the very young, very old and immunocompromised patients, so Trend data may represent a less healthy patient population and not necessarily general community prevalence. Conversely there are sites that perform a significant number of tests for the outpatient setting. This may create variability among the clinical sites’ percent positivity and introduces a challenge to comparing pathogen intensity between sites.

The uncertainties surrounding the testing algorithm and the precise patient population tested should not interfere with determining the onset, peak and duration of the pathogen season at each institution. These limitations on the data are likely to be common among almost all current surveillance systems for similar reasons. Given these concerns, the agreement between the percent positivity of Flu A/B as determined by Trend and the percent positivity reported by CDC FluView Influenza is striking (Figure 3), supporting the validity and utility of the Trend data.

A second source of concern in the Trend data set is a consequence of removal of sample identification such that we cannot directly determine whether the sample was from a patient or was a non-clinical sample (verification test, QC or PT) and should be removed from further epidemiological analysis. We estimate that non-patient testing makes up approximately 1.8% of the total FilmArray RP tests. Automated detection algorithms remove 3.5% of the total RP tests, including approximately half of the non-clinical samples. With the exception of the four positive tests, the clinical samples removed by filtering should be a random sampling of all patient tests. The remaining 1% fraction of non-patient tests have essentially no impact on the Trend evaluation of pathogen prevalence but they do make it more difficult to perform high resolution analysis of pathogen co-detections. This is especially true for co-detections of low prevalence organisms where QC positives are likely to be more common than real positives. Future updates to the FilmArray software will simplify the process by which the instrument operator can tag tests of non-patient samples, thereby largely eliminating the need to filter such test results from the Trend database before analysis.

### Trend Variety

The total positivity rate of the FilmArray RP test varies from a low of 38% in the summer months to a high of 75% in December and January, with a yearly average of approximately 60%. Figure 5A shows that the average number of different circulating pathogens at a single institution can vary from eight up to 11 during the winter months. Even during the peak periods of ILI, many respiratory infections are due to other viruses (Figure 2C) that can present clinically in a similar fashion [68, 69]. Therefore, the presumption of an influenza infection based on reported influenza percent positivity, without diagnostic testing for the virus, can lead to the inappropriate use of anti-viral agents [70]. Conversely, without comprehensive testing, a negative influenza or RSV test can lead to prescription of an unnecessary antibiotic. FilmArray Trend data can be a valuable aid for antimicrobial stewardship programs because it provides real-time information regarding the causes of respiratory infections and highlights the prevalence of viral infections.

As previously observed [66], the viruses that share the winter seasonality of influenza demonstrate annual or biennial behavior. It is possible that the viruses that share an influenza-like seasonality but do not show a two year cycle (RSV and hMPV) are actually alternating strains but the FilmArray RP Test does not detect this difference (e.g. the FilmArray RP does not differentiate between RSV A and RSV B). Adenovirus and the bacteria show constant occurrence through the year; HRV is in a unique class with peaks in the fall and spring.

Detection of multiple respiratory viruses in the same patient has been reported before. In the Trend dataset the rate of dual and triple co-detections was approximately 7.7 %, with HRV/EV as the organism most commonly observed in a co-detection. Some viruses, such as ADV and CoVs are detected in a mixed infection more than 50% of the time (Figure 4). In principle a FilmArray RP positive result may represent detection of residual pathogen nucleic acid from a previous infection that has resolved. However, several studies suggest that coinfections are associated with more severe disease ([71-73], see also discussion in [74]). In such cases, information about multiple detections can provide infection control practitioners with data that can assist in bed management and in the assessment of risk for nosocomial infections in a patient population that has been segregated by the occurrence of a common pathogen. Such information can prevent the introduction of a new pathogen associated with cohorting patients during busy respiratory seasons [75-77].

The question of whether different respiratory pathogens interfere with, or facilitate, growth in a human host is of some interest and not well understood. With the right data it can be studied at the population [78], individual [79], and cellular level [74]. Because the Trend data still includes some non-patient tests, we have chosen not to analyze every possible dual or triple infection individually. Rather we have taken a global approach and compared the overall rate of observed co-detections with MIE, which is a measure of the diversity of viruses circulating in a specific region and time period. MIE is similar, but not identical, to PIE (Probability of Interspecific Encounter [80]), also referred to as the Gini-Simpson index (1-D, where D is the Simpson s index [81, 82]), which is used in ecology as a measure of the species diversity of a region. Similarly, the circulating pathogen number of Figure 5A is identical to the Species Richness measure of ecology. We calculate MIE using frequencies (P_i_) of pathogen positivity per FilmArray test and note that the sum of all pathogen frequencies can add up to more than 100% because of co-detections or be less than 100% because of the presence of negative tests. In this regard, MIE differs from PIE because it is not a probability measure.

Figure 5B shows that the observed rate of co-detections is a constant fraction of MIE (approximately one quarter as indicated by the linear regression of Figure 5-Figure supplement 1).This observation suggests that, in the aggregate, respiratory pathogens are appearing in co-infections at a rate that can be predicted by their observed abundance. Data however may be biased by the patient population tested and the type of respiratory disease. The data also does not rule out that there are particular respiratory pathogens that occur more or less often in mixed infections than predicted by their individual percent positivity rates. [74, 83]. As we improve our ability to remove non-patient test results from the Trend dataset we will be able to characterize specific virus co-detection rates and their significance [65, 66, 78, 79, 84-86].

### Trend Value

As with weather forecasting, there is both a theoretical and a practical interest in predicting the next few weeks or months of the respiratory season [87-90]. FilmArray Trend contributes to infectious disease forecasting efforts because the data is timely and comprehensive. As the number of sites participating in Trend increases it will be possible to localize the reported infections to smaller geographical regions. At a high enough density of Trend sites, patterns of movement of respiratory pathogens across the US will become visible in a way that has not been easily observed before now.

The Trend RP data is summarized as a stacked graph that shows the percentage contribution of each pathogen to what is currently being detected by FilmArray RP testing (Figure 2B, www.syndromictrends.com). This analysis does not take into account changes in the rate of testing over a given season, information that should provide additional data regarding disease intensity and severity. In contrast the simple metric, Test Utilization Rate (TUR), describes the non-normalized rate of FilmArray test usage and serves as a surrogate for the level of syndromic disease that health care providers observe (Figure 2A).

TUR suffers from two defects. First, it is closely linked to the sales of the FilmArray test and thus is proprietary data that BioFire does not share (Google took a similar position in regard to releasing the search queries used by Google Flu Trends [14]). Second, TUR is driven by both the demand for testing and the growth in FilmArray product adoption and increasing acceptance and usage by health care providers. A useful step beyond TUR would be a normalized Test Utilization Rate that can adjust for the underlying growth of testing unrelated to the intensity and duration of the respiratory disease season. An increase in a normalized TUR metric may indicate the prevalence of circulating respiratory viruses and the intensity of respiratory disease overall. Likewise, an increase in the normalized metric, concomitant with an increase in negative tests, may indicate the occurrence of an outbreak caused by an emerging pathogen.

Public health agencies (PHAs), which include local and state health departments and the CDC, are specifically exempt under an HIPAA provision that allows clinical laboratories to disclose PHI to the PHA for specified public health purposes [91]. The exemption includes follow up studies on reportable infectious diseases. Real-time pathogen specific syndromic surveillance systems such as Trend will allow state PHAs to more rapidly identify, acquire, and test residual samples from potential outbreaks. Conversely, perceived “outbreaks” may actually be coincidental multi-organism seasonal surges, and rapid analysis by Trend-like systems could prevent timely and costly outbreak investigation.

Given the movement in healthcare technology towards greater vertical integration of a hospital’s data, the Bottom-Out approach exemplified by Trend will face more competition from Top-Out approaches (Figure 1, see for example GermWatch in Utah, [24]), because these systems can capture patient information (e.g. age, gender, and patient address) that is critical for more detailed epidemiological analysis. However, combining PHI with the diagnostic test result in the Top-Out approach makes these systems more complex and difficult to implement and may limit participation by health care institutions. Ironically, Bottom-Out data export systems have a role to play in the development of Top-Out systems because Bottom-Out export provides a rapid and efficient means to quality check the data flowing from Top-Out systems. FilmArray Trend data could also be combined with data derived from other automated diagnostic platforms [92, 93]. This work might best be accomplished by a third party that is viewed as independent and impartial. For example, in the case of data originating in the US, a federal institution or a private foundation could host a database to which IVD manufacturers would contribute their different syndromic test results. The benefits of a more complete view of circulating pathogens should outweigh the complexities of combining data from different platforms.

### Outlook

FilmArray Trend is a novel surveillance tool for simultaneously monitoring multiple syndromic diseases that has demonstrated promise in expanding our knowledge of the epidemiology of infectious diseases. Indeed, the close correlation of seasonal respiratory viruses tracked by Trend with reported CDC ILI highlights the major contributory role of multiple respiratory pathogens beyond influenza to ILI. The national and global expansion of Trend will provide a comprehensive tool to study the impact of co-infections, understand the role of previously underappreciated pathogens, and clarify true disease epidemiology. Finally, systems like Trend will be essential for the rapid identification of disease anomalies indicating potential emergent outbreaks, thereby providing an independent tool for public health surveillance.

## Methods

### FilmArray System and data output

The FilmArray System performs nucleic acid purification, reverse transcription, nested multiplex PCR amplification and DNA melt curve analysis for up to 30 targets in 63 minutes ([37], www.biofiredx.com). The FilmArray disposable pouch is prepared by injecting hydration solution and, separately, the patient sample, i.e., for the FilmArray RP test, a nasopharyngeal swab in viral transport media. The pouch is loaded into the FilmArray Instrument and the operator scans the barcode (containing panel type, lot and serial numbers) and then scans or manually enters information into the “Sample” field. Additional free form text can be entered into a “Tag” field. Software on the computer connected to the FilmArray Instrument analyses the amplification products and indicates each pathogen target detected. The results are presented as a PDF report and can be exported to the laboratory’s LIS. Information about the patient (gender, age, date of birth etc.) is not recorded by the instrument unless specifically entered.

As of 2016, more than 4,000 FilmArray instruments have been placed in clinical use worldwide. Approximately 80% of these systems are in the US. The FilmArray RP test is United States Food and Drug Administration (US FDA) designated as Clinical Laboratory Improvement Act (CLIA) moderate status (can be used in moderate to high complexity laboratories).

### HIPAA and the use of de identified patient data for epidemiological tracking

The Health Insurance Portability and Accountability Act of 1996 (HIPAA) and the Health Information Technology for Economic and Clinical Health Act (HITECH, a part of the American Recovery and Reinvestment Act of 2009) apply to the use of IVD test results. The US Department of Health and Human Services (DHHS) has issued regulations that set standards for the use of patient data (see links at [94]). Under HIPAA, laboratories that perform the FilmArray test on patient samples are “Covered Entities.” If laboratories send PHI to a third party (such as the IVD manufacturer) then that party is acting as a business associate (BA) of the Covered Entity and a Business Associate Agreement (BAA) would be required.

The HIPAA Privacy Rule [95] dictates that Covered Entities and, by extension, their BAs, may only disclose PHI with the patient’s written authorization for the purposes of treatment, payment or normal health care operations, and a small number of additional exemptions [96]. Trend software resolves HIPAA concerns by removing all PHI before the test results leave the Covered Entity. This makes it highly unlikely that BioFire, or a malicious intruder into the Trend database, could associate the FilmArray test results with a patient. Because PHI is not exported, BioFire is not a BA and a Data Use Agreement (DUA) between the laboratory and BioFire addresses the export of FilmArray data to the Trend database. Demonstrating that Trend database does not contain PHI has been critical to recruiting institutions to this project.

### Expert determination of de identification

The HIPAA Privacy Rule allows for release of patient data without prior authorization as long as it has been properly de-identified [97]. There are two acceptable routes to de-identification,1) the Safe Harbor approach wherein the data are stripped of an enumerated list of 18 identifiers classified as PHI and there is no indication (i.e., actual knowledge, that the remaining information would lead back to the individual) and 2) the Expert Determination approach wherein a person with experience in relevant statistical and scientific principles evaluates the PHI in conjunction with other reasonably available records, establishes a protocol for de-identification and certifies that the protocol allows only a small risk that PHI may be disclosed to an anticipated recipient.

The goal of the Trend project is to provide a near real-time view of the changes in pathogen prevalence; therefore, it is important to be able to retrieve the date when a FilmArray test is performed. However, retrieving the date conflicts with the Safe Harbor approach because the date of a test is PHI (the year of the test is not PHI but working with just the year defeats the purpose of tracking prevalence through the season). For this reason we followed the Expert Determination approach to manage data export.

The study took into consideration data that are available on participating clinical laboratory FilmArray Instruments, BioFire’s own customer database, the proposed Trend database and publicly available data sources. We analyzed how combinations of this information could be used by an adversary to identify an individual in the dataset thereby disclosing PHI [98]. The results of this study (summarized in Table 2) provided recommendations for development and site enrollment criteria for Trend, and for BioFire operating procedures.

In accord with the recommended actions, information in the fields that may be used to distinguish a patient is obfuscated (through truncation or binning) to ensure that a combination of these fields cannot be used to identify a specific patient [97]. For example, the time and date of the test are dynamically binned so that a minimum number of tests of one panel type are included in each bin prior to export. This ensures that a sufficient quantity of test results are uploaded to the database from one site at one time so that there is very low risk that patient identity can be inferred from knowledge of the start time of the test.

If an adversary were to infiltrate the safeguards of the database, and wished to know specific patient test results from a specific location on a given day, no unique records would exist. The combination of deleting the sample identification (ID) field, binning the test date range, and truncating the FilmArray pouch serial number ensures that the remaining information is never unique, which indicates that there is a low risk of misuse of data.

### FilmArray Trend Client Software and Database

The Trend client software resides on the computer associated with the FilmArray Instrument(s). The computer makes a secure, HTTPS, connection to services hosted by Amazon Web Services. Authenticated data submissions are stored in a database hosted by Amazon Relational Database Service. Both services have been configured to be HIPAA Security Rule compliant [99]. Trend client software requires that the computer has Internet access to make secure outbound web requests. The HTTPS protocol is industry standard technology used for secure banking and web applications. The Website software authenticates the Trend data before saving the exported de-identified data to the Trend Database.

### Sites used in the Pilot Trend project

The clinical laboratory sites participating in the pilot Trend project are shown in Table 3. The HHS regions are as defined by the CDC [5].

### Cleaning the Trend data

FilmArray Trend should display only valid test results acquired as part of the normal clinical testing of patients. Results from tests where an instrument, software or process control failure occurred have been removed from the data set. In addition, there are situations in which FilmArray customers test non-patient samples: FilmArray clinical customers can purchase Quality Control material from several different commercial sources and can make their own QC material by using cultured organism (from, for example, Zeptometrix, Buffalo, NY) either individually or in combination, or by using patient samples that previously tested positive for one or more organisms on the panel. Although BioFire provides recommendations for performing initial test validations [100] that involve combining pathogens in pools of 1-4 per test as well as negative controls, clinical laboratories may use other combinations of analytes. The signature of several of the commercial mixes can be identified simply by observing the set of organisms present. For example MMQC (Maine Molecular Quality Controls, Saco, ME) makes synthetic RNA mixes (M211 v1.1 and M212 v1.1) that explain 98% and 97% of the positive signals for seven and 12 analyte test results, respectively (Table 1, All, lines 7 and 12).

Additional QC testing is associated with specific events: a site adopts a new panel, qualifies an instrument returned from repair or, more frequently, qualifies a new lot of FilmArray RP pouches. Since we cannot be sure which tests are performed on control material, we identify events that are associated with test results containing more than three positive calls and flag all tests associated with these events for removal. Table 1-Figure Supplement 1 presents an example in the case of the QC tests performed to qualify a new lot of FilmArray RP pouches. In the first six days after a new lot of FilmArray pouches is received, tests with more than three positive organism calls make up 50, 48, 18, 10, 7, and 3% of the results. Thus, for a new lot of pouches, we remove the first three days of testing to minimize the contribution from control runs, thereby deleting 1.4% of all test results from Trend.

Clinical laboratories also perform periodic proficiency testing using test samples provided by either CAP (College of American Pathologists, Northfield IL) or API (American Proficiency Institute, Traverse City, MI). This testing occurs at defined time periods during the year and BioFire is notified. Thus, we have knowledge of the organism mixes being tested as well as the number of such tests. Because only 0.05% of the results in the Trend data set are predicted to be part of CAP or API proficiency testing we have not removed them from the present study.

Because all patient identifiers are stripped before data are exported, there is no automated means to detect repeat tests on the same patient. However, the level of this testing is less than 1% of the total due to test cost and the fact that repeat testing should not be used for test-of-cure. For these reasons, the occurrence of patient duplicate testing would be rare and would not impact the overall Trend dataset. At present we cannot easily remove test results that are part of research protocols. Users at the Trend Pilot sites are instructed to add the Tag: “No Trend” to such tests (at present 0.03% of the Trend dataset) but due to the manual nature of the process we do not know how well this request is observed in practice.

### Test Utilization Rate (TUR) and Pathogen Detection Rate

The FilmArray RP TUR metric is defined as the non-normalized number of RP patient test results generated each week on average across the Trend sites (computed as a centered three week moving average). To calculate the pathogen detection rate (as displayed in Figure 2A and on the Trend website) we compute the rate for each organism at each institution as a centered three week moving average. In order to adjust for the capacity differences between sites, a national aggregate is calculated as the unweighted average of individual site rates Only data from sites contributing more than 30 tests per week is included to avoid noise from small numbers of tests. Because the calculation of pathogen detection rate includes results from patients with multiple detections the detection rate for all organisms can, in theory, add up to greater than one. In practice this does not occur.

### Comparison to the CDC Influenza observed rate of detection

The CDC FluView rate of Flu A and Flu B detections as well as the reported incidence of ILI are taken from the CDC website [101]. Only the CDC data from the HHS regions that contained Trend Pilot sites were used for calculating the rate of influenza detections (Table 3).

### Calculation of co detection rates and related measures

Pathogen co-detections are defined as FilmArray tests in which two or three organisms are detected. We also calculated two other measures that relate to co-detections: the Circulating Pathogen Number and the Measure of Interspecific Encounter (MIE). Both of these time series measures are calculated for each site and week, a centered five-week moving average is computed and then an unweighted average of all sites is used to create a national aggregate. The five week moving average is used to reduce noise due to small numbers of samples within a week at some sites.

More specifically the Circulating Pathogen Number is simply the count of the unique organisms detected at a site during a one-week period. MIE is calculated from the frequencies of each organism at a site (number of positive test results for an organism divided by the number of FilmArray tests performed at that site). To reduce noise, we only include site data if more than 10 FilmArray tests were performed in that week. If P_1_ P_N_ are the percentage detection of the N different organisms circulating at a single site over a single week then MIE is defined as:

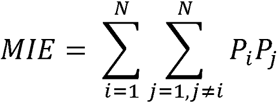

Conceptually, MIE is an attempt to estimate the likelihood that a patient infected with one organism may be infected with another unique organism circulating in the population at a given period in time, resulting in a co-infection.

## Additional information

### Acknowledgements

We acknowledge funding from NIH grant 5U01AI074419 (L.M., K.M.R. and M.A.P) and NIH/NIAID contract HHSN272201600002C (MAP). We thank Chris Thurston and Spencer Rose (BioFire Defense) for building the final Trend public website; Andrew Wallin (BioFire Defense) for reviewing the MIE data analysis; Anna Hoffee (BioFire Diagnostics) for assistance with the figures; Mark Pallansch (CDC), Kirsten St. George (New York State Department of Health) and Allyn Nakashima (Utah Department of Health) for useful discussions; Anne Blaschke and colleagues at BioFire Diagnostics and BioFire Defense for reviewing the manuscript.

### Author contributions

L.M., K.M.R. and M.A.P. developed the Trend concept. P.H.G. and C.C.G provided early input on the scope of the project. L.M. designed the DUAs, recruited the sites and provided overall coordination of the project. C.V.C. coordinated the clinical site software installations, L.M. and R.N. designed and supervised the coding of the data export software, database and web interface for the data. L.M. and A.N.F. performed the data analysis and prepared the figures. B.A.M. reviewed the expert determination studies. S.V.S., B.M.A. and J.D.J. reviewed the epidemiological analysis. F.S.N. tested the first release of the data export software. F.S.N., P. H. G., A.L., D.J., V.D, J.D.B., S.S., K.A.S., H.S., R.S., S.J., J.A.D., J.C.W., K.L., F.M., S.L.R, M.A-R., P.D.F., G.A.S., S.J.M., C.C.R., J.F.M. implemented the Trend export software at their respective institutions. C.C.G. and M.A.P. provided periodic review of the project. M.A.P., L. M. and C.C.G. wrote the manuscript. All authors reviewed the final manuscript.

### Author ORCIDs

Lindsay Meyers 0000-0003-2994-274X

Christine C. Ginocchio 0000-0002-8200-0324

Aimie N. Faucett 0000-0003-4590-4276

Frederick S. Nolte 0000-0002-4943-9220

Per H. Gesteland 0000-0002-9869-8122

Diane Janowiak 0000-0002-9208-0647

Virginia Donovan 0000-0002-9296-6931

Jennifer Dien Bard 0000-0003-0524-9473

Silvia Spitzer 0000-0002-2642-4771

Kathleen A. Stellrecht 0000-0002-0160-6034

Hossein Salimnia 0000-0002-8069-5185

Rangaraj Selvarangan 0000-0003-3275-6657

Stefan Juretschko 0000-0002-4658-6276

Judy A. Daly 0000-0003-4801-7023

Kristy Lindsey 0000-0002-8172-5964

Franklin Moore 0000-0002-3019-0437

Sharon L. Reed 0000-0001-7615-9366

Paul D. Fey 0000-0003-0939-6884

Camille V. Cook 0000-0001-6071-4898

Jay D. Jones 0000-0001-9091-5316

Samuel V. Scarpino 0000-0001-5716-2770

Benjamin M. Althouse 0000-0002-5464-654X

Kirk M. Ririe 0000-0003-0620-7419

Bradley A. Malin 0000-0003-3040-5175

Mark A. Poritz 0000-0002-5060-0560

### Competing interests

L.M., A.N.F., R.K.N., C.V.C, J.D.J., K.M.R., C.C.G. and M.A.P. are present or former employees of bioMérieux, Inc. or its subsidiaries. bioMérieux markets the FilmArray System and Trend. F.S.N., P.H.G., D.J., V.D., A.L., J.D.B., S.S., K.A.S., H.S., R.S., S.J., J.A.D., J.C.W.,

K.L., F.M., S.L.R., M.A-R., P.D.F., G.A.S., S.J.M., S.V.S., B.M.A., are research contractors of BioFire Diagnostics for the development of the FilmArray Trend system. CCR and JFM are members of the Trend Working Group. B.A.M. is a paid consultant of BioFire Diagnostics.

## Supplementary Figures

**Table 1-Figure Supplement 1:**
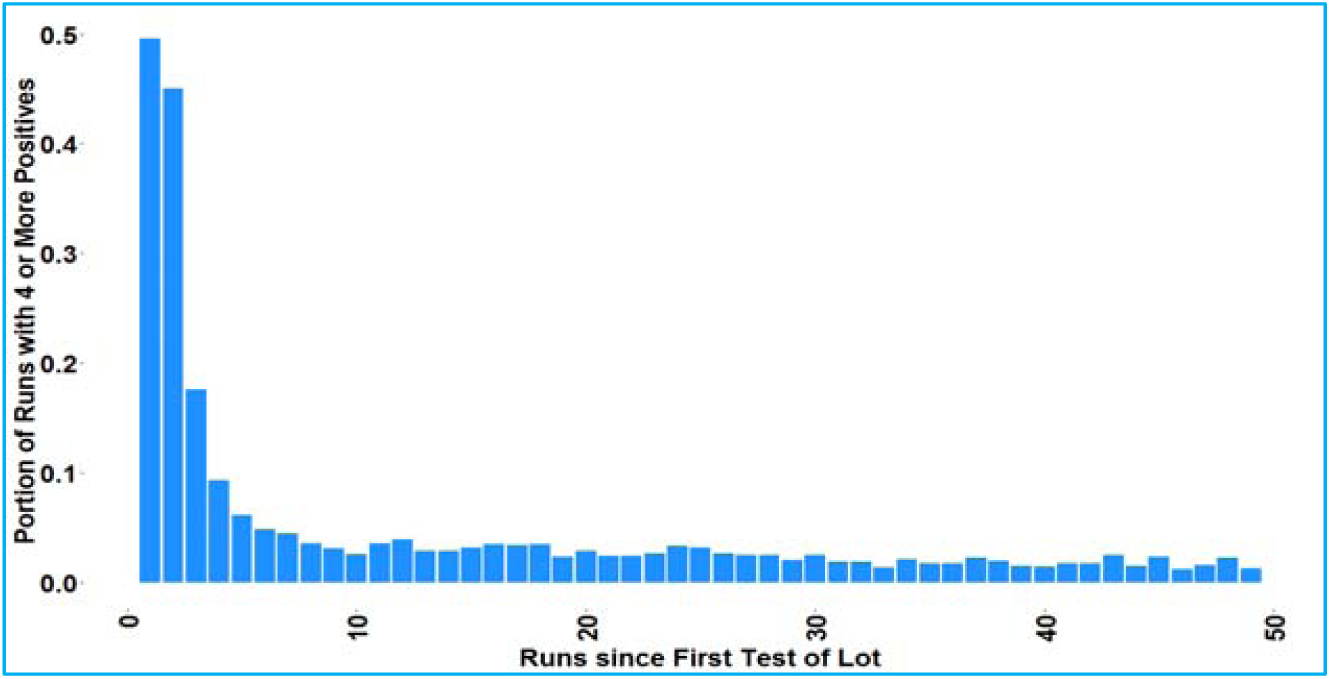
QC runs are performed when a new lot of FilmArray RP pouches is introduced. Histogram of count of FilmArray RP tests with more than three positive results after a new lot of FilmArray RP pouches is introduced at a clinical site. The X-axis is the sequential number of RP tests associated with the new lot that have been performed at that site.

**Figure 2-Figure Supplement 1:**
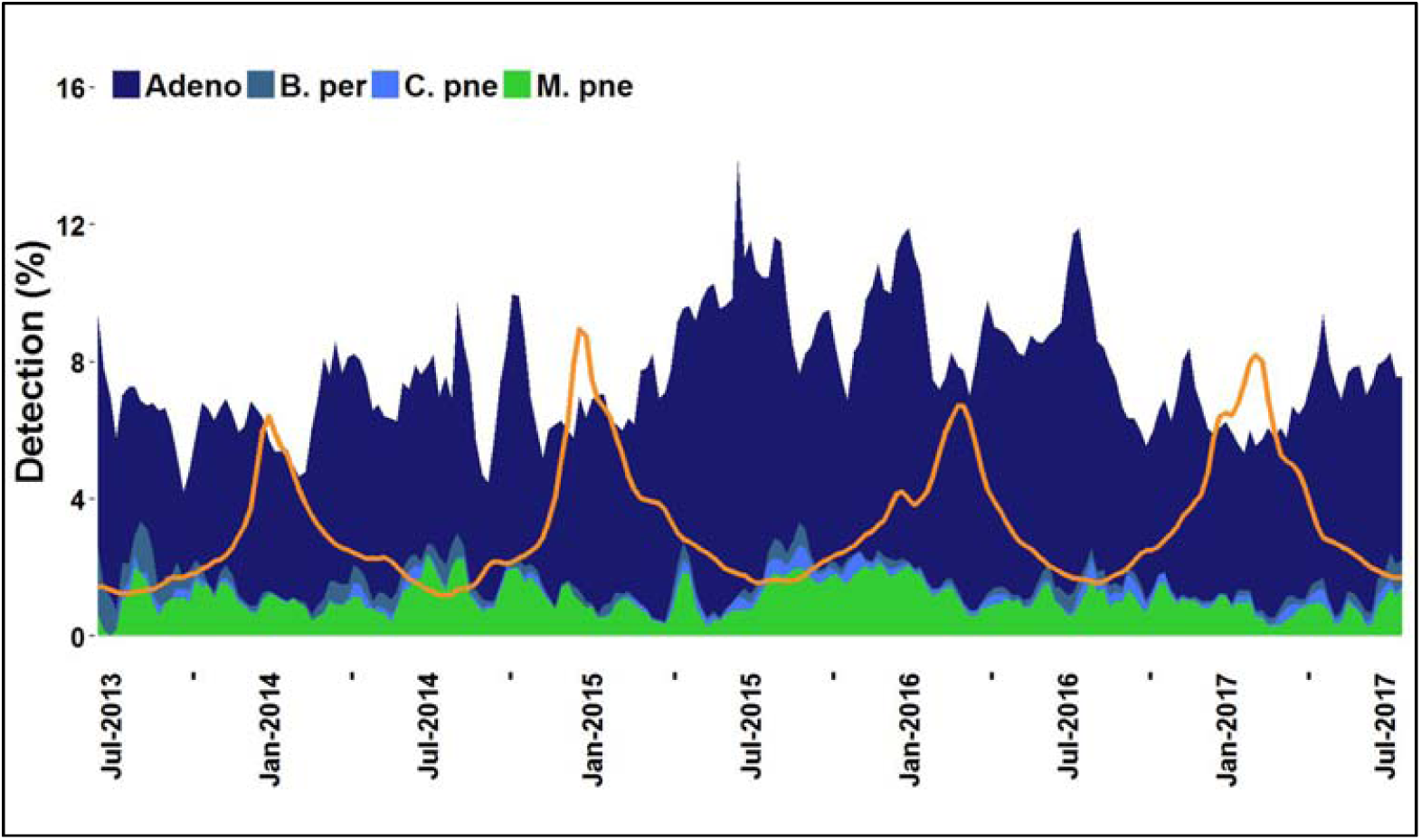
Detection of adenovirus and three bacteria. Detection of adenovirus and the three bacterial pathogens (*B. pertussis, C. pneumoniae*, and *M. pneumoniae*) in the Trend dataset displayed as a stacked area graph. CDC measured ILI is indicated (orange line).

**Figure 2-Figure Supplement 2:**
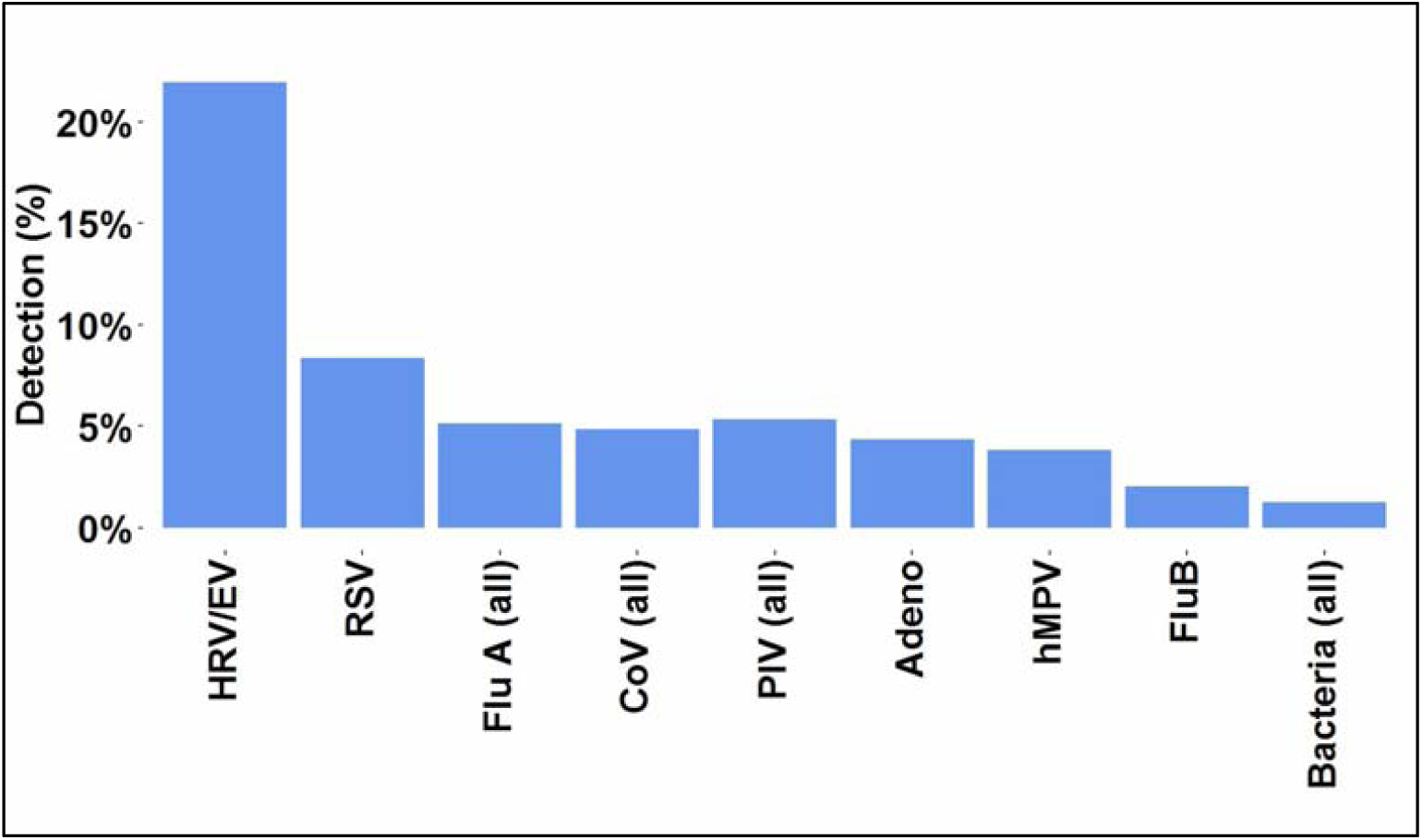
Detection of FilmArray RP organisms by type. Percent detection rates aggregated from all participating Trend sites. The percent positive detection per test for each organism group is ordered by abundance.

**Figure 2-Figure Supplement 3:**
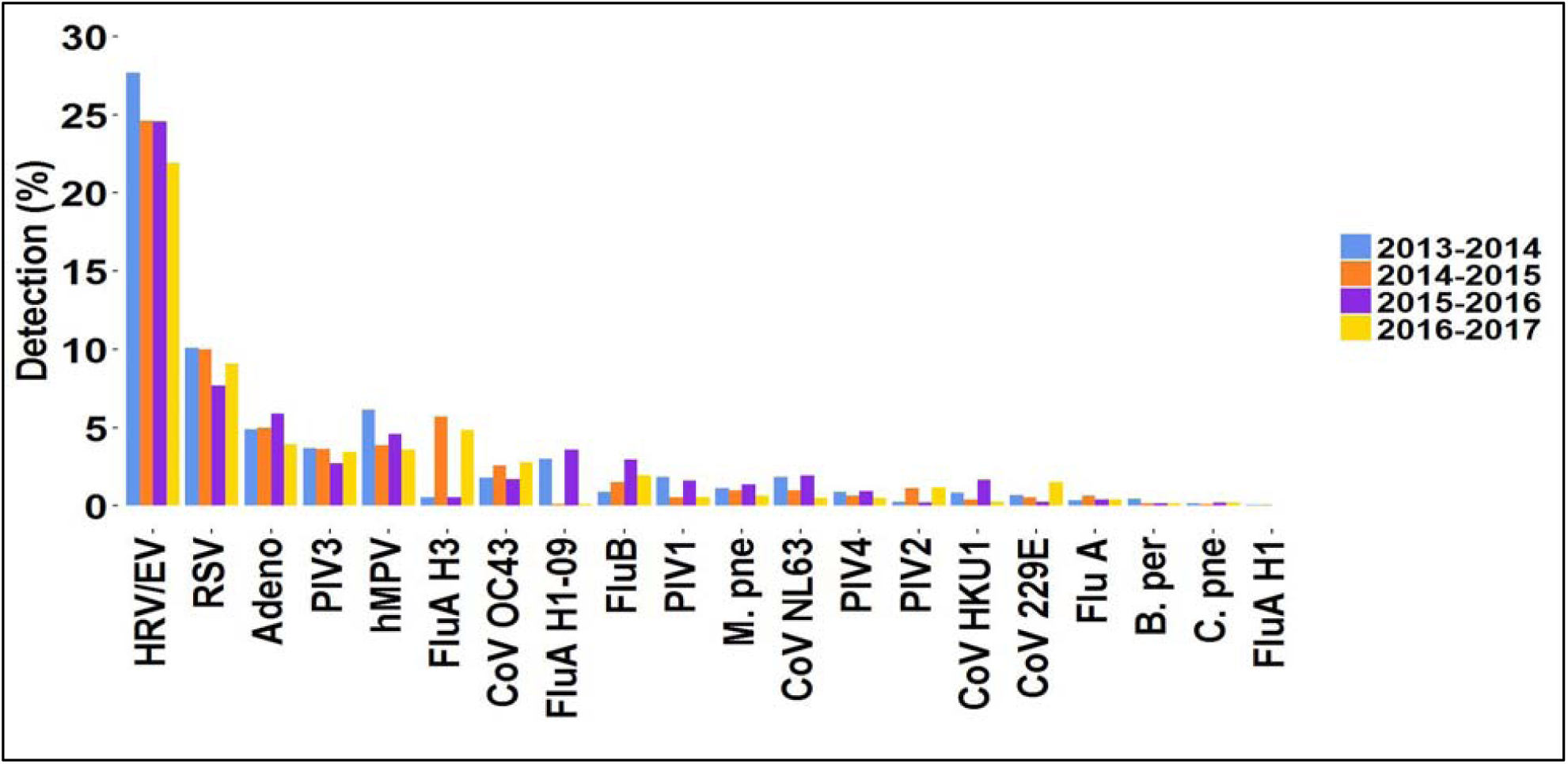
Organism detection by type and year. Percent detection rates from all participating Trend sites for each FilmArray RP organism are broken out by respiratory year. Organisms are ordered by total abundance across the four years.

**Figure 5-Figure Supplement 1:**
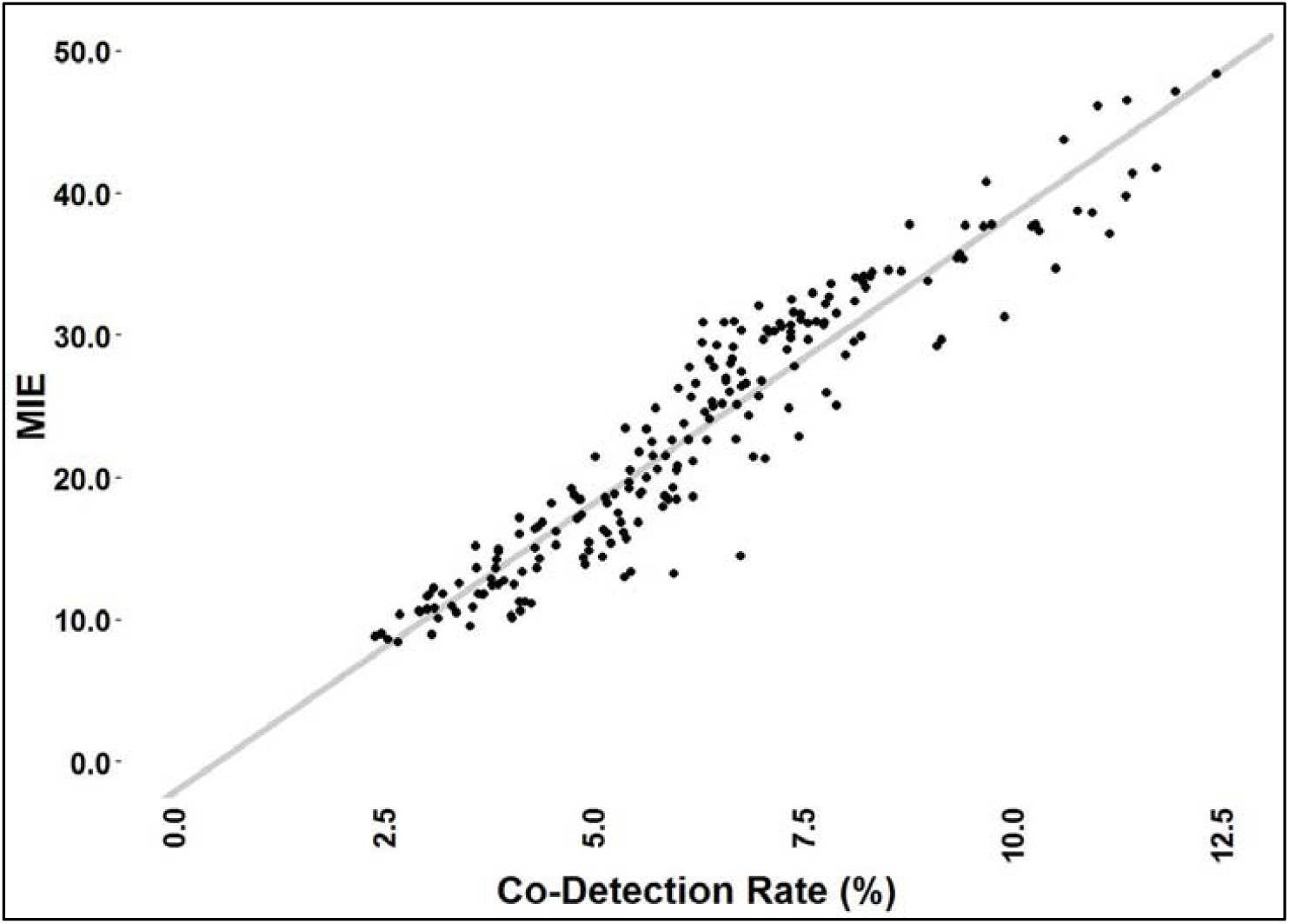
Linear regression of MIE and observed co detections. The time series data in figure 5B shown as a scatter plot. The equation of the linear regression is: MIE = 4.05153* Co-Detection rate-0.0206 with an R^2^ value of 0.9003

